# The genome of *Talinum fruticosum*

**DOI:** 10.1101/2023.04.20.537669

**Authors:** Dominik Brilhaus, Alisandra K. Denton, Eva Maleckova, Vanessa Reichel-Deland, Andreas P. M. Weber

## Abstract

Research on crassulacean acid metabolism (CAM) has in recent years focused on obligate CAM species, such as *Kalanchoë fedtschenkoi* and pineapple (*Ananas comosus*). To fully understand the plasticity of the CAM pathway, its evolutionary trajectory and regulation, genomic resources of additional species, including facultative CAM species are desirable. To this end, we sequenced the genome and full-length transcripts (Iso-Seq) of the facultative CAM dicot *Talinum fruticosum*.

The provided resources may aid in CAM engineering as an approach to improving crop water-use efficiency.

## INTRODUCTION

Crassulacean acid metabolism (CAM) has frequently and convergently evolved as a carbon-concentrating mechanism in vascular plants. It is the most water-use efficient photosynthetic strategy, with some CAM species requiring as little as one sixth of the water budget of C3 species (Yang et al., 2015a). The key towards reduced water demands as compared to C_3_ and C_4_ species is the rescheduling of atmospheric CO2 fixation to the dark period and thus temporal separation of primary and secondary carbon assimilation (Fig. 1). In the dark, stomata open and atmospheric CO2 is fixed in the form of HCO3-by PHOSPHO*ENOL*PYRUVATE CARBOXYLASE (PEPC), yielding oxaloacetate, which is immediately reduced to malate by MALATE DEHYDROGENASE (MDH) and then stored in the vacuole as malic acid. During the following light period, stomata close and nocturnally accumulated malic acid is released from the vacuole and then decarboxylated enzymatically to provide CO2 for the secondary fixation catalysed by RIBULOSE-1,5-BISPHOSPHATE CARBOXYLASE/OXIDASE (RuBisCO). At dawn and dusk, two transitory phases occur: the switch from PEPC-to RuBisCO-mediated carbon fixation at the dark/light transition, and re-opening of stomata towards the end of the light period to enable CO2 assimilation by RuBisCO as malic acid reserves decline (Osmond, 1978; Lüttge, 2002; Borland et al., 2009).

**Figure 1.**
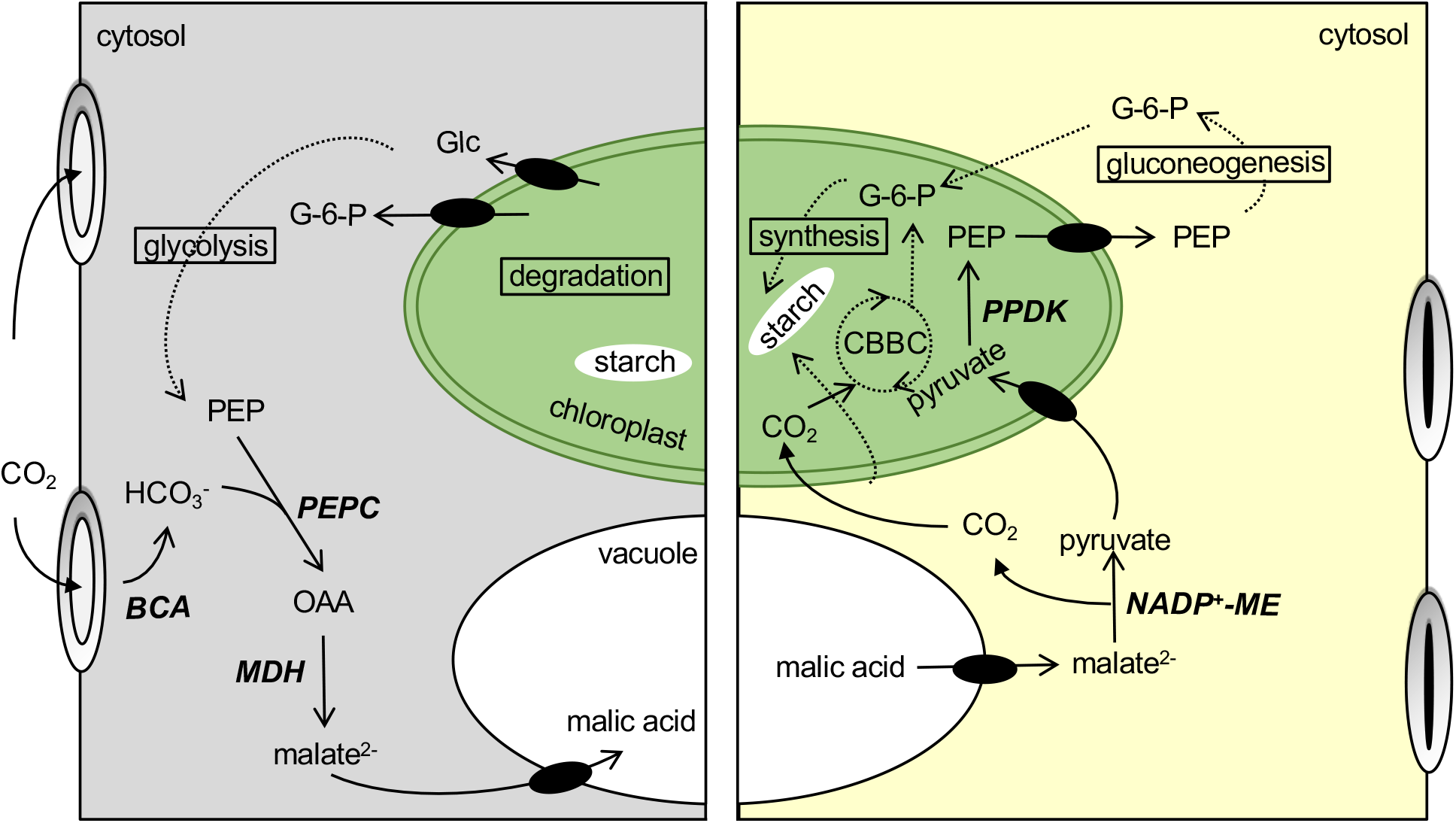
A simplified scheme of the CAM pathway. The key enzymatic conversion of both the light and the dark phase are shown. Enzyme names are shown in bold and italics, black ovals represent known or putative membrane transporters and dashed lines stand for multi-step enzymatic conversions with only substrates and products shown. BCA, β-carbonic anhydrase; CBBC, Calvin-Benson-Basshman cycle; Glc, glucose; G-6-P, glucose-6-phosphate; MDH, malate dehydrogenase; NADP^+^-ME, NADP^+^-dependent malic enzyme; OAA, oxaloacetate; PEP, phosphoenolpyruvate; PEPC, phosphoenolpyruvate carboxylase; PPDK, pyruvate phosphate dikinase.

CAM is a highly plastic adaptation to low water availability and relatively few species rely exclusively on carbon assimilation via this pathway. A striking example of this plasticity are so-called facultative CAM plants, which mostly rely on C_3_ assimilation and only upon exposure to stress, such as drought, switch to CAM mode. Unlike the ontogenetically pre-programmed C_3_-to-CAM transition in obligate CAM species (*e*.*g. Ananas comosus, Phalaenopsis equestris* and *Kalanchoë fedtschenkoi*), facultative CAM is reversible upon stress relieve (Wai et al., 2019; Winter, 2019a; Winter and Holtum, 2014). *Mesembryanthemum crystallinum* is the best described facultative CAM species, but more cases of facultative CAM have been described with more of the plant diversity being explored. Other facultative CAM species include *Talinum fruticosum* (formerly *T. triangulare*) and several C_4_-CAM *Portulaca* species (Brilhaus et al., 2016; Maleckova et al., 2019; Winter, 2019b). Facultative CAM also evolved in monocots (*e*.*g*. the orchid *Dendrobium catenatum*) and in the fern *Vittaria lineata* (Minardi et al., 2014; Zhang et al., 2016). Facultative CAM species present powerful experimental models to identify crucial components of the CAM pathway, such as CAM-specific enzyme isoforms and how they differ from their C_3_ variants. These models further allow investigating regulatory mechanisms underlying CAM induction and reversion to the C_3_ mode of carbon assimilation (Brilhaus et al., 2016; Maleckova et al., 2019; Winter and Holtum, 2014).

The flexible transition between fast growth in the C_3_ mode and inducible drought resilience, makes engineering facultative CAM a promising strategy for crop improvement (Schiller and Bräutigam, 2021). Successful CAM engineering efforts depend on detailed understanding of the regulatory mechanisms controlling CAM induction and how re-programming of carbon assimilation is synchronised with other metabolic processes (Liu et al., 2018a; Borland et al., 2014; Yang et al., 2015a). The evolution of CAM likely occurred via amino acid substitutions affecting kinetic, regulatory and binding properties of CAM proteins as well as altered timing of gene expression, frequently accompanied by changes in maximal abundance of gene products (Silvera et al., 2010; Cushman et al., 2008; Yang et al., 2017).

A detailed description of CAM-enabling evolutionary changes and the shared evolutionary trajectory between different plant lineages remain limited by the availability of (facultative) CAM plant genomes. The present work provides the first genome sequence of a truly facultative, *i*.*e*. fully C_3_-reversible, CAM dicotyledonous species.

## RESULTS

### Assembly of the *Talinum fruticosum* genome

*T. fruticosum* genomic DNA was sequenced using Nanopore and Illumina sequencing technologies to benefit from sequencing of long fragments and low error rates, respectively. The genome was assembled from 8,125,273 Nanopore reads with Flye (Kolmogorov et al., 2019a) and reaching a mean coverage of 141x (Tab. 1). It was polished with 273,931,287 paired-end (PE) Illumina reads using POLCA (Zimin and Salzberg, 2020). The assembly size was 621.7 Mbp (Tab. 1). It consists of 344 contigs with an N50 of nearly 6.7 Mbp (Tab. 1). The presence of 98% of highly conserved *Embryophyta* genes, as determined by Busco (Simão et al., 2015), indicated a highly complete assembly in genic regions. Repetitive sequences accounted for 397,312,403 bp (64% of the assembly). The most abundant repeats were long terminal repeat (LTR) retrotransposons, with Copia and Gypsy elements accounting for 21.5% and 9.2% of the genome assembly, respectively. Besides the nuclear genome, the chloroplast genome was obtained relying on a reference-based assembly process. The plastome sequence of *T. paniculatum* was used as a reference to map PE reads with GetOrganelle v1.7.1 (Jin et al., 2020). With 156,811 bp in size and 85 protein-coding genes, the resulting assembly was highly similar to the used reference sequence (Liu et al., 2018b).

**Table 1.** Statistics of genome assembly for *Talinum fruticosum*.

| Parameter | Value |
| --- | --- |
| Total assembly length | 621,663,653 bp |
| Number of contigs | 344 |
| N50 | 6,679,480 bp |
| L50 | 33 |

### Genome annotation

Structural gene annotations were generated using Helixer (Stiehler et al., 2020, Holst et al., 2023) using the hybrid convolutional and bidirectional long-short term memory model ‘HybridModel’. The specific trained instance of the model was ‘land_plant_v0.3_m_0100’. Default parameters were used except as otherwise mentioned below: the subsequence length was increased to 106920 bp. During inference, overlapping was activated using a core-length of 53460 bp and an offset of 13365 bp. Phase prediction was used. The obtained annotation was manually boosted using RNA-seq and Iso-seq data.

### The phylogenetic placement of *Talinum fruticosum*

The phylogeny of *T. fruticosum* was analysed with OrthoFinder (Emms and Kelly, 2019). Protein sequences of primary (*i*.*e*., longest) isoforms obtained with Helixer were compared to protein sequences of 29 species that represent the phylogenetic diversity of *Embryophyta* and include a diversity of photosynthetic types. In the resulting species tree, *T. fruticosum* was placed in the “ACPT” clade of *Caryophyllales* together with two *Portulaca* species (Fig. 2).

**Figure 2.**
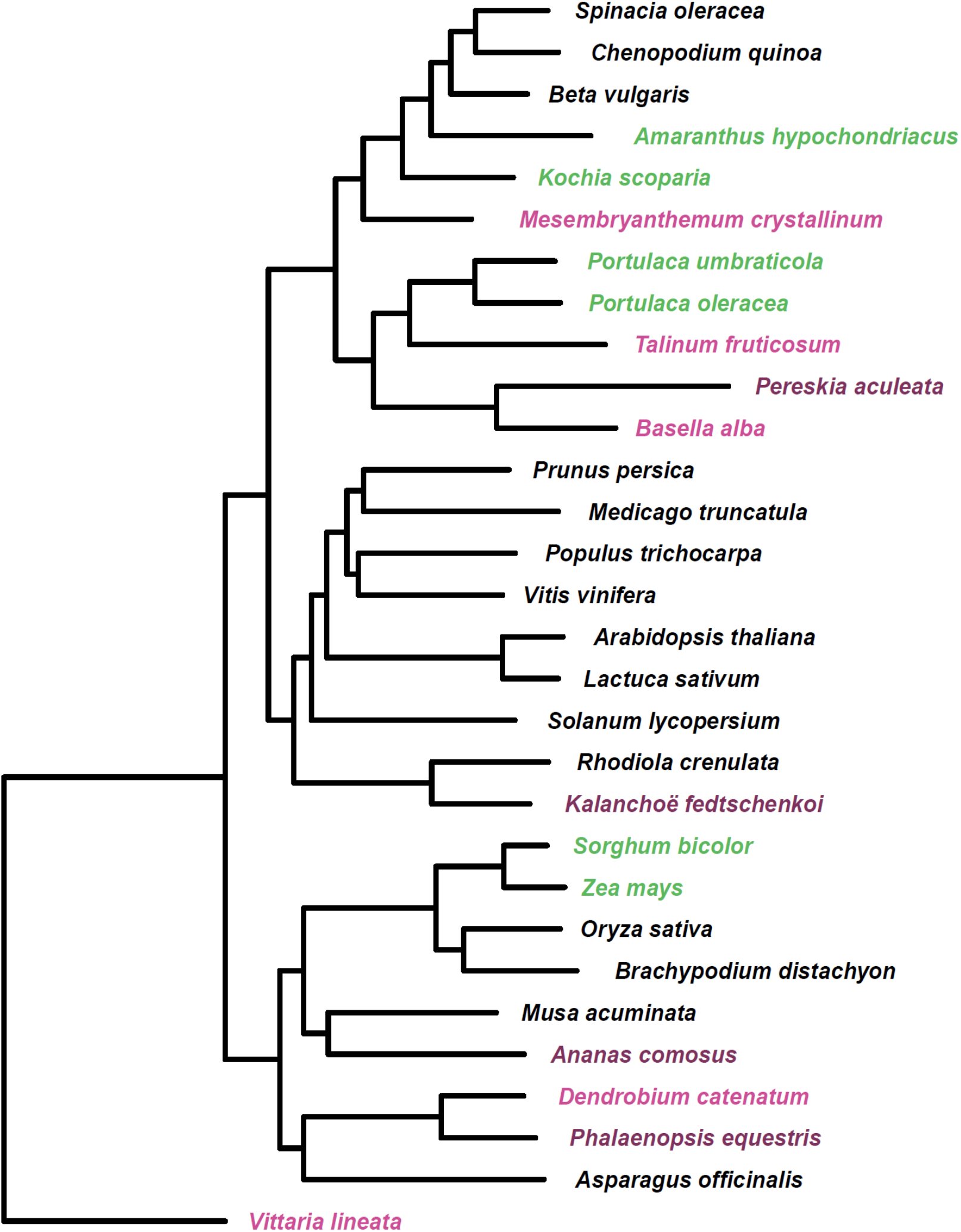
Phylogenetic placement of *Talinum fruticosum* based on STAG algorithm incorporated in OrthoFinder (Emms and Kelly, 2019).

### A CAM-specific *PEPC* isoform

The major PEPC orthogroup (OG0001398) comprised six proteins of *T. fruticosum*, the majority of them showing high sequence homology to Arabidopsis PEPC2 protein. The individual *T. fruticosum* PEPC isoforms differed not only in their amino acid sequence but also in diurnal pattern of transcript abundance in response to both drought and exogenous ABA. We propose *Tf_contig_062_000131* as the CAM-specific isoform. *Tf_contig_062_000131* transcripts accumulated rapidly in response to exogenous ABA (Fig. 3B), peaked at the light-dark transition under fully induced CAM (*i*.*e*. 12 days of water withdrawal), and diminished after re-watering (Fig. 3B).

**Figure 3.**
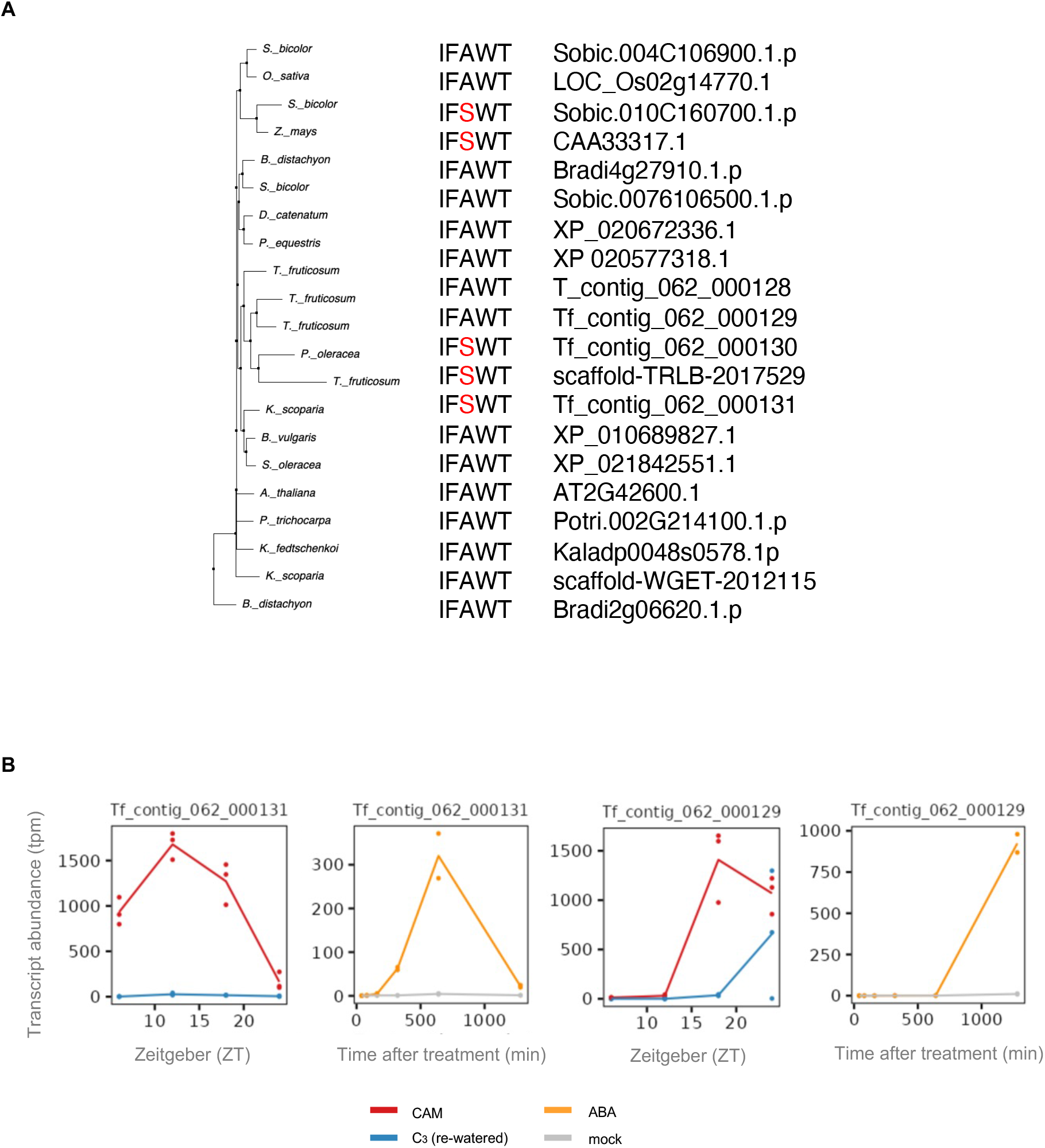
Identification of putative CAM-isoforms of phospho*enol*pyruvate carboxylase. (A) Multiple sequence alignment (MAFFT) of deduced protein sequences of Arabidopsis phosphoenolpyruvate carboxylase (PEPC2) orthologs. For simplicity, only residues associated with post-translation regulation and determining kinetic properties are shown. (B) Transcript abundance patterns of two most abundant, ABA- and drought-inducible genes.

The *Tf_contig_062_000131* protein contains a regulatory serine residue at position 780 (numbering based on PEP1 of *Z. mays* – UniProt ID: CAA33317), which is typical of C4 PEPC proteins and not observed in C_3_ PEPCs (Fig. 3A). This distinguishes *Tf_contig_062_000131* from other *T. fruticosum* PEPC isoforms, including the likewise inducible and highly abundant isoform *Tf_contig_062_000129* (Fig. 3A), but also from *PEPC2* orthologs of other CAM species, including *M. crystallinum, A. comosus* and *K. fedtschenkoi* (Fig. 3A).

## DISCUSSION

(Facultative) CAM engineering has been proposed as a strategy to improve the water-use efficiency of crops. Its success however depends on detailed understanding of regulatory mechanisms underlying CAM, which can be well studied in facultative CAM species. Here, we present the genome sequence of *T. fruticosum*.

Using Nanopore sequencing together with PE Illumina sequencing, we assembled a genome with mean coverages of 141 and 84, respectively (Tab. 1). The total assembly size was 621.7 Mbp.Based on protein sequence similarity, *T. fruticosum* was placed in the “ACPT” clade of *Caryophyllales* and in near proximity of C3 relatives *B. vulgaris* and *S. oleracea* (Fig. 2), confirming the current phylogeny of *Caryophyllales* (Yang et al., 2018, 2015b). The presence of a vast majority (98.7%) of highly conserved *Embryophyta* orthologs in the assembly indicates high completeness, which makes it a valuable resource for (facultative) CAM research.

Duplications of whole genomes or individual genes are a major driving force for evolution (Kanazawa et al., 2009). This was also reported for the evolution of CAM photosynthesis, for example in *K. fedtschenkoi* where a whole genome duplication is associated with diversified diel expression patterns between the gene copies of *e*.*g*., *PEPC* and *PEPCK* (Yang et al., 2017). In contrast, sequencing of *P. equestris* and *A. comosus* genomes did not reveal associations between recent whole genome duplications and CAM evolution (D’Andrea et al., 2015; Ming et al., 2015). This likely holds true also for *T. fruticosum* since there was no general trend towards expansion of the examined CAM-related orthogroups unlike in triploid *Musa acuminata*. Prominent exceptions were *PEPC* and *PPCK* orthogroups (*i*.*e*., enzymes involved in nocturnal carbon assimilation) which showed increased number of predicted gene models compared to the C_3_ species *A. thaliana, B. vulgaris*, and *S. oleracea* as well as CAM monocots included in the analysis.

We hypothesize that diversification of protein-coding and regulatory sequences are – at least in the facultative CAM species *T. fruticosum –* not mutually exclusive as illustrated with the example of *Tf_contig_062_000131 (PEPC2)*, which we propose as a CAM-specific isoform. It is not the only isoform showing altered transcript abundance in response to exogenous ABA, prolonged drought and subsequent re-watering, but its transcripts accumulated to the highest level on the analysed time points in both the ABA and the drought experiments. Further its predicted protein sequence features a Ser residue at position 780 (numbering based PEP1 of *Z. mays*) (Fig. 3). Replacement of the ancestral Ala residue that is typical of C3 species by a phosphorylatable Ser residue lowers the enzyme’s affinity to PEP, allowing substrate accumulation to higher levels (Bläsing et al., 2000). This substitution is a well-described phenomenon in C_4_ species, such as *Flaveria trinervia* and C_4_ grasses (Bläsing et al., 2000; Christin et al., 2007), but less frequent among CAM species ((Christin et al., 2014) and Fig. 3).

However, an amino acid change from Arg, His or Lys residues to Asp can still be associated with increased PEPC activity as shown for *K. fedtschenkoi* and *P. equestris* (Yang et al., 2017). This however is also not found in other CAM species such as pineapple or *Agavoideae* members, suggesting that this mutation is not a necessary requirement for the CAM pathway (Heyduk et al., 2022). It can rather be assumed that there is variability of the pathway, meaning it could be essential to have one PEPC copy that needs to be recruited (Heyduk et al., 2022). Therefore, we are suggesting Tf_contig_062_000131 to be the CAM-enabling PEPC copy in *T. fruticosum*.

Comparative genomics of a diversity of CAM species and their C_3_ relatives will enable identification of changes in protein-coding and regulatory sequences. It will become feasible to understand crucial evolutionary trajectories towards CAM-enabling changes as well as to identify changes specific to certain lineages only with possibly more fine-tuning functions.

It is expected that besides core CAM enzymes, additional protein activities must be regulated accordingly, such as enzymes of carbohydrate metabolism and membrane transporters (Chen et al., 2020). It remains to be investigated whether transport proteins of CAM species differ from their C3 homologs. This work demonstrates the importance of availability of genomic resources for diverse (facultative) CAM species. Utilizing those, it will be possible to identify potential CAM-enabling changes, such as shared temporal patterns of gene expression of core CAM genes and regulators of the CAM pathway, expansion of certain gene families or altered kinetic properties. With follow-up experiments that might include generation of gene regulatory networks and its respective loss-of-function mutants, promoter-transcription factor (TF) binding assays as well as studying TF-TF interactions, it will be possible to identify the most suitable building blocks for CAM engineering, as previously proposed by Yang et al., 2015.

## METHODS

### Plant Material and Growth Conditions

For genome and transcriptome sequencing, seeds of *Talinum fruticosum* obtained after five to six subsequent, controlled self-pollination events were germinated in D 400 soil with Cocopor (Stender) and grown in a controlled-environment plant chamber (MobyLux GroBanks, CLF Plant Climatics) under the following conditions: 12 h light/12 h dark at 25 °C/23 °C. The light intensity at the leaf level was 150–200 µmol s-1 m-2. For isoform sequencing, a batch of seeds was sterilised by washing twice in 70% ethanol for 15 min followed by two washes with sterile water (15 min each) and germinated on ½ Murashige and Skoog medium with 3% sucrose to obtain sterile root tissue.

### DNA Extraction and Nanopore Sequencing

Progeny after five selfing events was used for Nanopore sequencing and grown under the above-described conditions. At the age of 25 days after sowing (DAS), the plants were kept at constant darkness for 36 hours to remove starch. Afterwards, whole shoots were harvested, snap-frozen in liquid nitrogen and stored at −80 °C until used. Approximately 10 g of snap frozen shoots were ground in liquid nitrogen to fine powder which was transferred to 200 ml homogenization buffer (HB) (10 mM Tris-Cl pH 9.5, 80 mM KCl, 100 mM EDTA, 1 mM spermidine, 1 mM spermine, 17.1% (w/v) sucrose). β-mercaptoethanol was added to a final concentration of 0.15% (v/v) and the sample was stirred for 10 min on ice. Subsequently, it was filtered through two layers of cheesecloth and one layer of miracloth, 5 ml 20% Triton X-100 (in HB) were added to the filtrate and stirred gently for additional 20 min on ice. Obtained lysate was centrifuged at 2,500 *g* for 20 min at 4 °C, the pellet was resuspended in 30 ml HB and filtered through two layers of miracloth. Afterwards, two more washing steps without any further filtering were performed.

Purified pelleted nuclei were resuspended in 2 ml G2 buffer (800 mM guanidine hydrochloride, 30 mM EDTA, 30 mM Tris base, 5% (v/v) Tween 20, 0.5% (v/v) Triton X-100), 4 µl RNase A [10 mg/ml] were added and incubated at 37 °C for 30 min. Afterwards, 45 µl proteinase K [20 mg/ml] were added and incubated at 50 °C for two hours, while occasionally inverting the tube. After spinning down at 5,000 *g* for 10 min at 4 °C, the supernatant was loaded on equilibrated Genomic-tip 20/G column (QIAGEN) and washed and eluted according to manufacturer’s instructions. DNA was precipitated with 0.7 volume of isopropanol and centrifuged at 5,000 *g* for 15 min at 4 °C. The pellet was washed two times in 500 µl cold 70% ethanol, each time followed by centrifugation at 5,000 *g* for 10 min at 4 °C. Upon discarding ethanol from the last washing step, the DNA pellet was air dried, 50 µl 10 mM Tris pH 8.5 were added to the pellet and DNA was allowed to resuspend by shaking overnight at 4 °C. Size and integrity of gDNA was examined with Fragment Analyzer™ (Advanced Analytical Technologies GmbH) and quantified with dsDNA HS Qubit assay, in both cases following the manufacturer’s protocol. Libraries were prepared from 1.5 µg gDNA with SQK-LSK109 Ligation Sequencing Kit (Oxford Nanopore Technologies) according to the manufacturer’s instructions and sequenced on the GridION™ X5 and the PromethION™ 24 platforms with R9.4.1 flow cells following the manufacturer’s guidelines. On-instrument base-calling was carried out by Guppy v3.2.8 and v3.2.10 (Oxford Nanopore Technologies).

### DNA Extraction and Illumina Sequencing

Progeny after six selfing events, which was grown under the above-described conditions, was used for PE Illumina sequencing. At the age of 30 DAS, plants were kept in the dark and harvested as for Nanopore sequencing.

DNA extraction was performed in aliquots of about 20 mg ground material. Wizard® Genomic DNA Purification Kit (Promega) was used and tissue lysis performed according to the manufacturer’s instructions. Subsequently, DNA precipitation was performed with 0.1 volume 3 M NaAc pH 5.2 and 0.7 volume room temperature isopropanol. After centrifugation at 5,600 *g* for 30 min at 4 °C, DNA pellets were washed twice with ice cold 70% ethanol and air dried prior to resuspending in 40 µl 10 mM Tris pH 8.5. Any remaining RNA was removed with 2 µl RNase A (Promega) added to a pool of DNA samples. After incubation at 37 °C for 30 minutes, DNA was purified with AMPure XP beads (Beckman Coulter) used in 1:1 ratio and eluted in 90 µl low TE buffer (10 mM Tris-HCl pH 8.0, 0.1 mM EDTA pH 8.0). DNA integrity was evaluated with a Fragment Analyzer (Advanced Analytical Technologies) using the HS NGS Fragment Kit (DNF-467, Agilent). dsDNA HS assay (Thermo Fisher Scientific) for Qubit™ was used for DNA quantification.

A sequencing library was prepared from 650 ng DNA, which was first fragmented to 350 bp using Covaris M220 (settings: duty cycle 20%, power 50 W, duration 65 s, temperature 20 °C.). VAHTS Universal DNA Library Prep Kit for Illumina V3 (version 8.1) was used to prepare the sequencing library. Manufacturer’s instructions were obeyed except for no final amplification was performed. The resulting library was assessed with Fragment Analyzer (Advanced Analytical Technologies) using the Genomic DNA Kit DNF-467 and quantified with KAPA Library Quantification Kit (Roche). A single lane of HiSeq3000 was used for sequencing with 2×151 bp read length in presence of 1% PhiX (Illumina). A total of 296,241,916 reads was obtained.

### Genome Assembly and Polishing

Prior to assembly, raw Nanopore reads labelled as “pass” were aligned to the chloroplast sequence of *T. fruticosum* assembled in this work and to a mitochondrial sequence of *Beta vulgaris* (NC_002511.2; (Kubo et al., 2000)) with minimap2 v2.17-r954-dirty (Li, 2018) and only Nanopore reads without a significant alignment to either organellar sequence (93.8% of all “pass” reads) were used for *de novo* assembly of the nuclear genome. Flye v2.7b-b1528 (Kolmogorov et al., 2019b) with a genome size estimation of 1 Gbp was used to obtain a draft assembly. This was subsequently polished with 273,931,287 PE reads retained after trimming (see Trimming of PE reads). Polishing was performed using POLCA, a tool available within the MaSuRCA v3.4.1 distribution (Zimin and Salzberg, 2020). Assembly statistics was obtained with QUAST v5.0.2 (Gurevich et al., 2013).

### Contaminant Removal

The polished genome assembly was aligned against the non-redundant nucleotide NCBI database (downloaded on 19.8.2019) using BLAST+ v2.6.0 (Camacho et al., 2009). The results were viewed in MEGAN v6.8.15 (Huson et al., 2016). Significant No hits outside *Viridiplantae* were obtained, but there were several contigs with no hits in the NCBI database. These were retained in the assembly.

### Genome Completeness Assessment

Genome completeness was determined using Benchmarking Universal Single-Copy Orthologs (BUSCO) v3.0.1 (Simão et al., 2015) with embryophyte_odb10 database of single-copy orthologs and default settings.

### Transcript Mapping

Full-length transcripts (PacBio sequencing) and previously generated PE mRNA-seq reads (Brilhaus et al., 2016) were used to generate extrinsic evidence for subsequent gene calling. Unique isoforms obtained with the isoeq3 pipeline and cDNA Cupcake scripts were mapped to the genome of *T. fruticosum* with GMAP v2017-05-03 (Wu and Watanabe, 2005) and sorted alignments were obtained with SAMtools v1.3.1 (Li et al., 2009). After trimming with Trimmomatic v 0.33, PE RNA-seq reads were mapped against the genome of *T. fruticosum* as well using HISAT2 v2.1.0 (Kim et al., 2015).

### Functional Annotation of the Nuclear Genome

For functional annotation, predicted protein sequences of *T. fruticosum* were compared to protein-coding sequences of 29 species using OrthoFinder. For species with reference genomes available, the primary transcript sequence sets were used. In the remaining cases, entire shotgun transcriptome assemblies were used without any filtering. Conclusions on protein function were based on presence of Arabidopsis proteins in the respective orthogroups. Besides that, typical protein signatures and domains were identified with InterProScan v5.51-85.0 (Jones et al., 2014) to confirm predictions based on Arabidopsis orthologs and gain insight on potentially novel genes.

### Trimming of Illumina Reads

Trimmomatic v0.33 (Bolger et al., 2014) was used to remove adapter sequences and low-quality bases from the raw PE reads and to keep only reads longer than 36 bases. The same quality and length criteria were applied to both WGS reads obtained both in the scope of whole genome sequencing and in both RNA-seq experiments (see Re-used data section).

### Mapping of RNA-seq Reads and Transcript Abundance Analyses

RNA-seq reads from both the ABA experiment (Maleckova et al., 2019) and the drought/re-watering experiment (Brilhaus et al., 2016) were mapped to the CDSs obtained with the above-described gene calling pipeline using Kallisto v0.45.1 (Bray et al., 2016).

### Accession Numbers

The sequencing reads generated in the course of this work are deposited in the Sequence Read Archive (SRA) of the National Center for Biotechnology Information (NCBI): Nanopore reads are available under accessions SRX9314398 and SRX9314397, Illumina PE reads are available as an accession SRX9210518, and Iso-Seq reads as an accession SRX9007794. The mRNA-seq reads re-used in this work (See also the Re-used data section of Materials and Methods) are available from the NCBI’s Gene Expression Omnibus (GEO) database under the following accessions: GSE116590 (ABA experiment; (Maleckova et al., 2019)), GSE70601 (drought/re-watering experiment; (Brilhaus et al., 2016)) and GSE99186 (additional reads from the same drought/re-watering experiment as in GSE70601). The nuclear genome assembly generated and used in this work is available under accession JAITYX000000000.1 (https://www.ncbi.nlm.nih.gov/nuccore/JAITYX000000000.1), the chloroplast genome under GenBank (NCBI) accession OK573457.1.

## Abbreviations

CAM: crassulacean acid metabolism
DAS: days after sowing
LTR: long terminal repeat
ORF: open reading frame
PE: paired-end
PEPC: phosphoenolpyruvate carboxylase

## Acknowledgements

Computational support and infrastructure were provided by the “Centre for Information and Media Technology” (ZIM) at the University of Düsseldorf (Germany). DNA and RNA-sequencing was performed by the Genomics and Transcriptomics Laboratory (GTL) of the West German Genome Center (WGCC) Düsseldorf. This work was funded by CEPLAS (Cluster of Excellence on Plant Science) supported by Deutsche Forschungsgemeinschaft within the Excellence Initiative (EXC 1028) and under Germany’s Excellence Strategy (EXC 2048/1 – Project ID: 390686111), by International Graduate Program for Plant Science (iGRAD-Plant; IGK 2466) and by a graduate fellowship by the German Academic Exchange Service (DAAD) to EM.

## Author Contributions

E.M. generated the homozygous lines, performed nucleic acid extractions, generated the assemblies, performed the downstream analyses and drafted the manuscript. D.B., A.K.D., V.R.-D. did bioinformatic analysis and wrote the manuscript. A.P.M.W. acquired financial support, provided the overall direction of the project and contributed to manuscript writing.

